# Adaptation and exogenous attention interact in the early visual cortex: A TMS study

**DOI:** 10.1101/2023.10.27.563093

**Authors:** Hsing-Hao Lee, Antonio Fernández, Marisa Carrasco

**Affiliations:** Department of Psychology, New York University, New York, NY 10003, USA; Center for Neural Sciences, New York University, New York, NY 10003, USA

**Keywords:** adaptation, contrast sensitivity, covert attention, transcranial magnetic stimulation (TMS), visual cortex

## Abstract

Transcranial magnetic stimulation (TMS) to early visual cortex modulates the effect of adaptation and eliminates the effect of exogenous (involuntary) attention on contrast sensitivity. Here we investigated whether adaptation modulates exogenous attention under TMS to V1/V2. Observers performed an orientation discrimination task while attending to one of two stimuli, with or without adaptation. Following an attentional cue, two stimuli were presented in the stimulated region and its contralateral symmetric region. A response cue indicated the stimulus whose orientation observers had to discriminate. Without adaptation, in the distractor-stimulated condition, contrast sensitivity increased at the attended location and decreased at the unattended location via response gain–but these effects were eliminated in the target-stimulated condition. Critically, after adaptation, exogenous attention altered performance similarly in both distractor-stimulated and target-stimulated conditions. These results reveal that (1) adaptation and attention interact in the early visual cortex, and (2) adaptation shields exogenous attention from TMS effects.

## Introduction

Due to the brain’s limited metabolic resources and the high energy cost of cortical computation, we are unable to process all the information available in the environment. To maximize perceptual performance, energy must be allocated according to task demands. Both visual adaptation and attention help manage the limited energy, optimizing visual processing and sensitivity^1-3^. On the one hand, adaptation reduces the visual system’s response to repetitive stimuli while enhancing sensitivity to non-adapted stimulus features^4^; e.g., prolonged viewing of a stimulus recenters contrast sensitivity away from the adaptor. Adaptation reduces sensitivity via contrast gain—the contrast response function (CRF) shifts rightward: higher stimulus contrast is required for observers to reach the same performance than before adaptation (**Figure 1A**)^5,6^. On the other hand, covert spatial attention–the selective processing of information at a specific location without shifting our gaze–enhances contrast sensitivity at the attended location and impairs it at unattended locations, via a “push-pull” mechanism^7,8^. Because orientation discriminability is contingent upon contrast sensitivity, we discriminate stimulus orientation better when covert attention is allocated to the stimulus location than elsewhere^2^.

**Figure 1.**
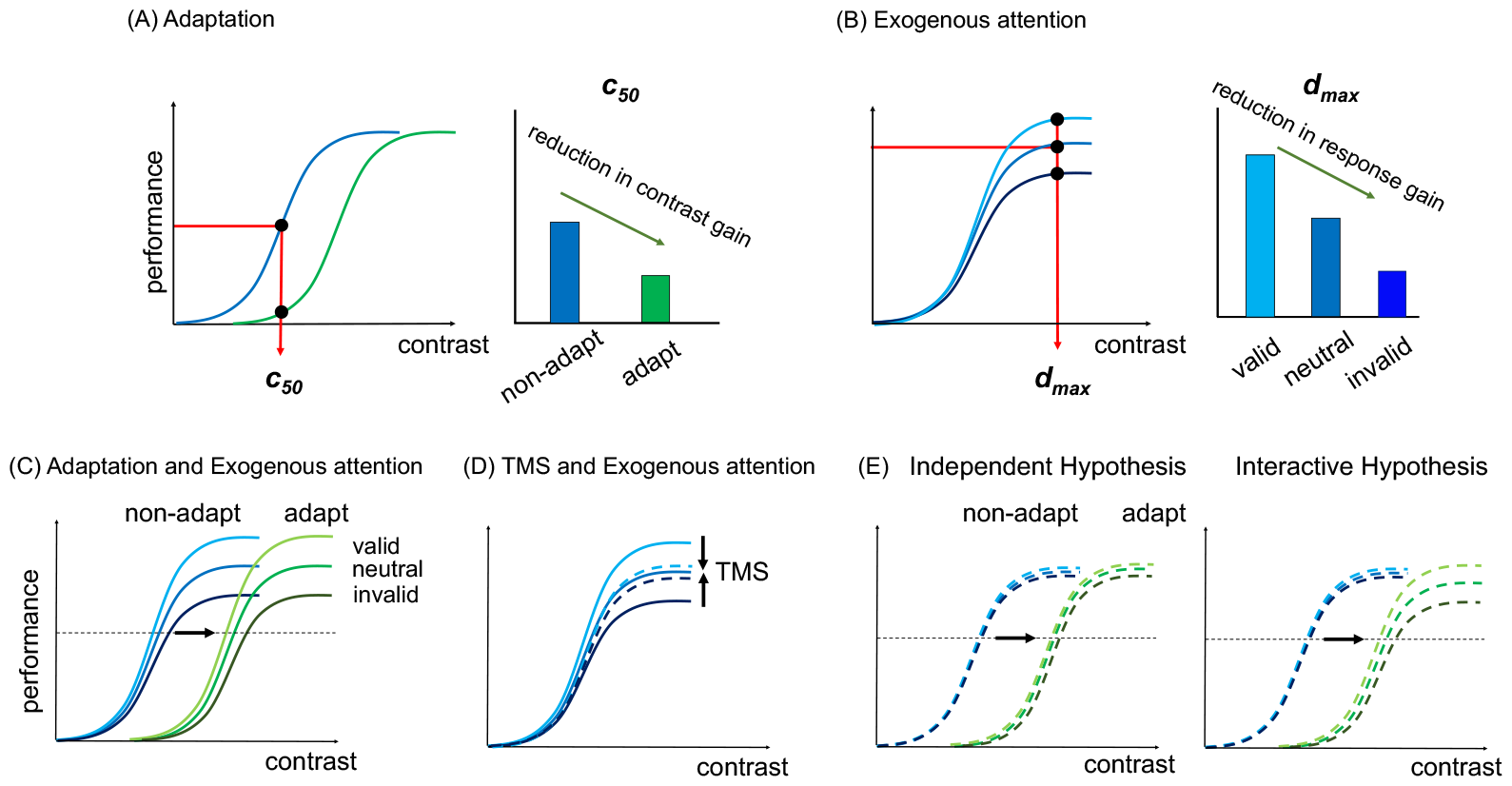
Effects of adaptation and attention effects on contrast sensitivity. (**A**) Adaptation reduces contrast gain: The c_50_ (semi-saturation point) is higher in the adapted than non-adapted condition^6,11,18,23^. (**B**) Exogenous attention modulates performance via response gain: performance at d’ max (asymptote) is highest in the valid, followed by neutral and invalid trials^2,7,10,11,33^. (**C**) Exogenous attention restores contrast sensitivity via response gain even if adaptation depresses overall contrast sensitivity via contrast gain^11^. (**D**) TMS to the target will disrupt the response gain brought by exogenous attention^10^. (**E**) Hypotheses: If the effects of adaptation and attention are independent in the early visual cortex, we should observe that the attentional effect is still eliminated by TMS after adaptation (left panel). Otherwise, the attentional effect will still emerge under the influence of TMS after adaptation (right panel).

There are two types of covert spatial attention: Endogenous attention is voluntary, conceptually driven and sustained; exogenous attention is involuntary, stimulus-driven and transient^2,9^. Exogenous attention primarily alters the CRF via response gain: an increase at the attended location and a decrease at the unattended location in the upper asymptote (**Figure 1B**)^7,10^. When jointly manipulated, adaptation and attention affect contrast sensitivity: while adaptation decreases contrast sensitivity via contrast gain, exogenous attention still alters sensitivity via response gain (**Figure 1C**)^11^.

Transcranial magnetic stimulation (TMS) induces a magnetic field that alters the local electric field in the brain^12-14^, thus enabling inferences regarding the causal role of specific brain areas to perception and cognition^15^. Effects of TMS on performance depend on TMS-pulse intensity^16^ and brain state^17^: When the initial state of the neuronal population is active, TMS suppresses activity, but when the initial state is suppressed, TMS disinhibits the neuronal population in adaptation^18,19^, covert attention^10,20^ and presaccadic attention^21^ studies.

TMS on early visual cortex (V1/V2) extinguishes the benefit and cost of exogenous attention on contrast sensitivity (**Figure 1D**)^10^. Observers were instructed to perform an orientation discrimination task while two TMS pulses were applied during stimulus presentation. A response cue indicated the patch whose orientation observers had to discriminate. The response cue either matched—target stimulated—or did not match—distractor stimulated—the stimulated side. When the distractor was stimulated, exogenous attention yielded the typical performance benefit and cost in the valid and invalid cue conditions, respectively, consistent with the exogenous attention effect without TMS^7,11^. But when the target was stimulated, all three conditions had similar performance (**Figure 1D**). By suppressing activity at the attended location and disinhibiting suppressed activity at the unattended location, this study provides evidence for the TMS state-dependent effect.

Adaptation^18,19^, exogenous attention^10^ and TMS^22^ each alter activity in early visual cortex. Here, we investigated whether adaptation and attention are independent or interactive by examining if TMS eliminates exogenous attentional effects after adaptation. Were these independent processes, adaptation would not modulate the effect of TMS on exogenous attention–TMS would still extinguish attentional benefits and costs. Were these interactive processes, adaptation would modulate the effect of TMS on exogenous attention (**Figure 1E**).

## Results

We titrated the tilt angle needed to achieve 75% correct discrimination performance and derive the semi-saturation point (c_50_; **Figure 1A**) of the CRF for each individual. The asymptote level was set at 80% contrast for all observers (d_max_; **Figure 1B**; see **Methods**). Thirteen observers discriminated whether a stimulus was tilted counterclockwise or clockwise off vertical at c_50_ or d_max_ contrasts. We tested observers’ performance at these two contrasts across attention, adaptation, and TMS conditions to infer contrast gain and response gain mechanisms.

The adaptation and non-adaptation sessions were administered on different days. In the adaptation sessions, observers experienced flickering Gabors (100% contrast) followed by an attentional cue and the stimuli. In the non-adaptation sessions, the procedure was the same but without the flickering Gabors (**Figure 2A**). Observers received two TMS pulses separated by 50 ms during target presentation (**Figure 2B)**. We presented the two stimuli in the TMS stimulated region and its symmetric location in the other hemifield; the stimulus was presented for each observer according to their phosphene location (**Figure 2C**). In half of the trials, the response cue instructed observers to report the orientation of the stimulus at the stimulated region (contralateral to TMS; <u>target-stimulated</u>), and in the other half, the symmetric region (ipsilateral; <u>distractor-stimulated</u>). In each case, the test was either preceded by a valid, invalid or neutral cue, with equal probability (**Figure 2A**; see **Methods**).

**Figure 2.**
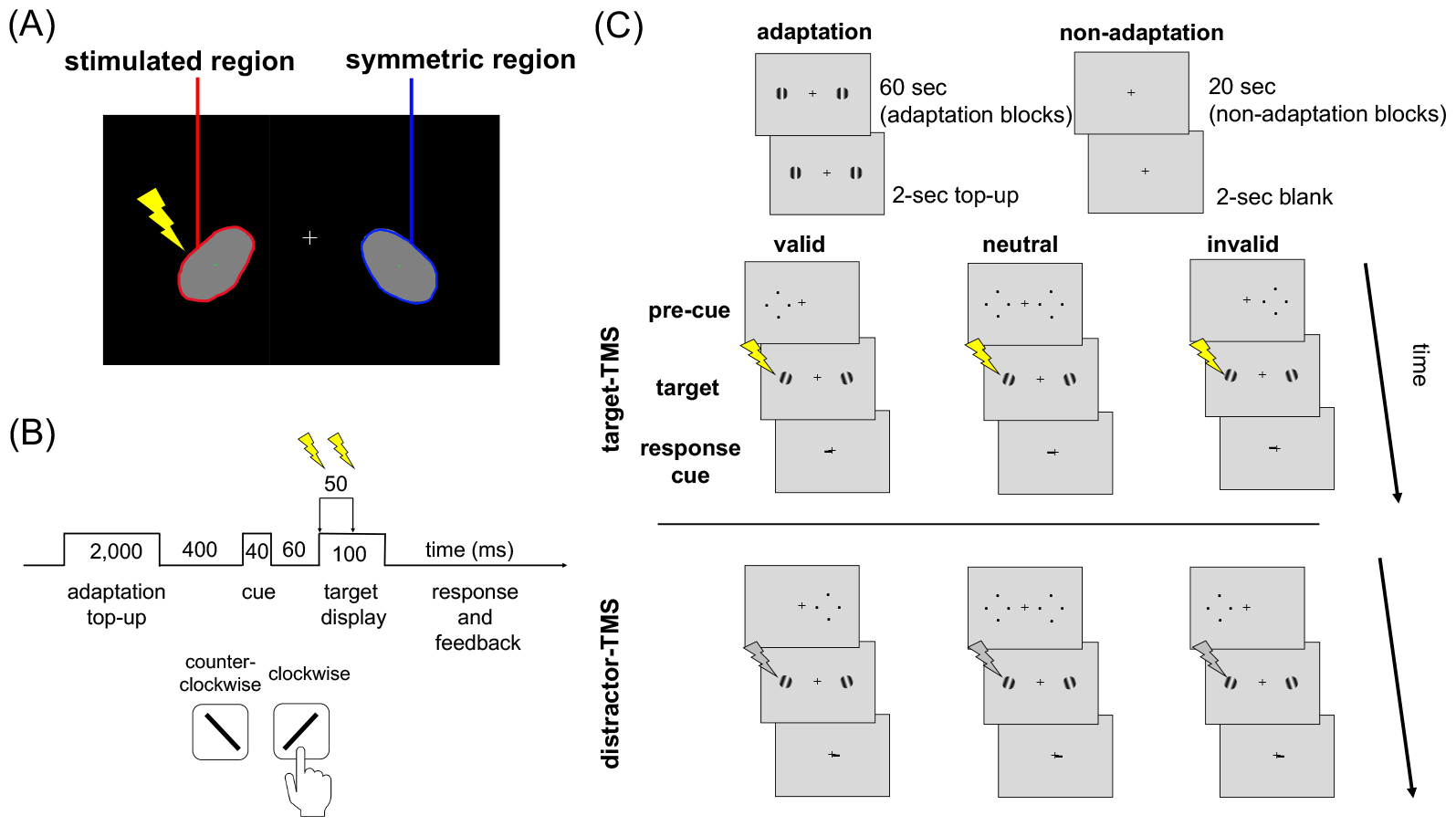
The psychophysics-TMS task. (**A**) Phosphene mapping: observers were stimulated near the occipital pole before they started the psychophysics-TMS task. They were instructed to draw the perceived phosphene outline using the cursor. This phosphene mapping procedure was repeated at the beginning of every session. (**B**) Trial timeline. Two TMS pulses were given during target presentation (separated by 50 ms). (**C**) The experimental design: Observers performed the adaptation or non-adaptation blocks in different experimental sessions. In the valid trial, the peripheral cue matched the location of the response cue. In the invalid trial, the peripheral cue mis-matched the location of the response cue. In the neutral trial, the peripheral cues were shown on both sides. In the target-TMS condition (middle panel), the response cue indicated the target in the stimulated region. In the distractor-TMS condition (bottom panel), the response cue indicated the target in the non-stimulated region (and the distractor was stimulated).

### Adaptation effects

To examine the adaptation effect on contrast sensitivity, we first assessed performance in the distractor-stimulated, neutral condition (**Figure 3)** –in which we expected no effect of TMS. A 2 (adaptation, non-adaptation) × 2 (c_50_, d_max_) within-subject analysis of variance (ANOVA) revealed higher performance (*d’)* in the d_max_ than c_50_ conditions [*F*(1,12)= 95.69, *p*<.001, *η*^*2*^=0.89) and an interaction [*F*(1,12)=11.81, *p*=.005, *η*^*2*^=0.5): Performance was lower at c_50_ contrast, [*t*(12)=4.02, *p*=.002, *d*=1.3], but not at the d_max_ contrast, [*t*(12)=2.01, *p*=.068], in the adaptation than non-adaptation conditions. This finding is consistent with adaptation depressing contrast sensitivity primarily via contrast gain^6,11,18,23^.

**Figure 3.**
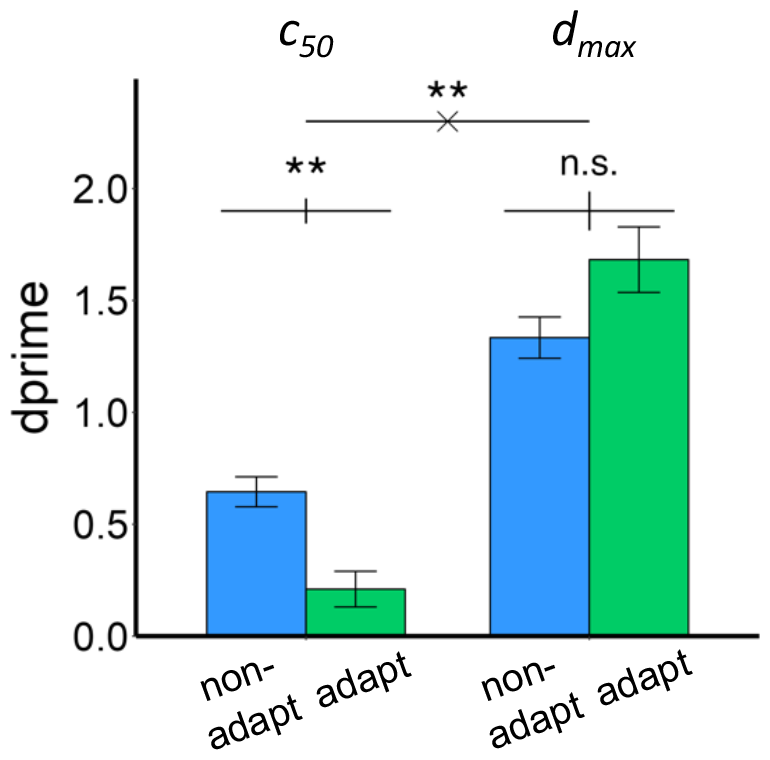
Performance (indexed by *d’*) in the distractor-stimulated neutral condition. An interaction revealed a lower *d’* in the c_50_ condition but not in the d_max_ condition. The error bars indicate ±1 SEM. * *p* < .01, * *p* < .05.

To explore the effects of attention under different adaptation conditions, we conducted a 4-way within-subject ANOVA on attention (valid, neutral, invalid), adaptation (adapt, non-adapt), TMS (distractor-, target-stimulated), and contrast (c_50_, d_max_). There were main effects of attention [*F*(2,24)=10.3, *p*<.001, *η*^*2*^=0.46], adaptation [*F*(1,12)=23.8, *p*<.001, *η*^*2*^=0.66] and contrast [*F*(1,12)=184.2, *p*<.001, *η*^*2*^=0.94]. There was no 4-way interaction [*F*(2,24)<1], but there were 3-way interactions among attention, adaptation and TMS [*F*(2,24) =5.9, *p*=.008, *η*^*2*^=0.33] and attention, adaptation and contrast [*F*(2,24)=3.96, *p*=.033, *η*^*2*^=0.25]. To interpret these 3-way interactions, we assessed the attention effect under TMS for c_50_ and d_max_ without and with adaptation.

### The attentional effects under TMS without adaptation

Overall, *d’* were lower than in the study of Fernández and Carrasco^10^. This difference resulted from the tilt and contrast stimulus parameters we used, based on pilot data so that attention and adaptation would have room to increase or decrease performance. We examined the attentional effect without adaptation using within-subjects ANOVAs on attention (valid, neutral, invalid) and TMS (distractor-TMS, target-TMS). For c_50_ (**Figure 4A**), there were no main effects on attention [*F*(2,24)=1.91, *p*=.169] or TMS [*F*(1,12)<1], nor was there an interaction [*F*(2,24)=1.75, *p*=.195].

**Figure 4.**
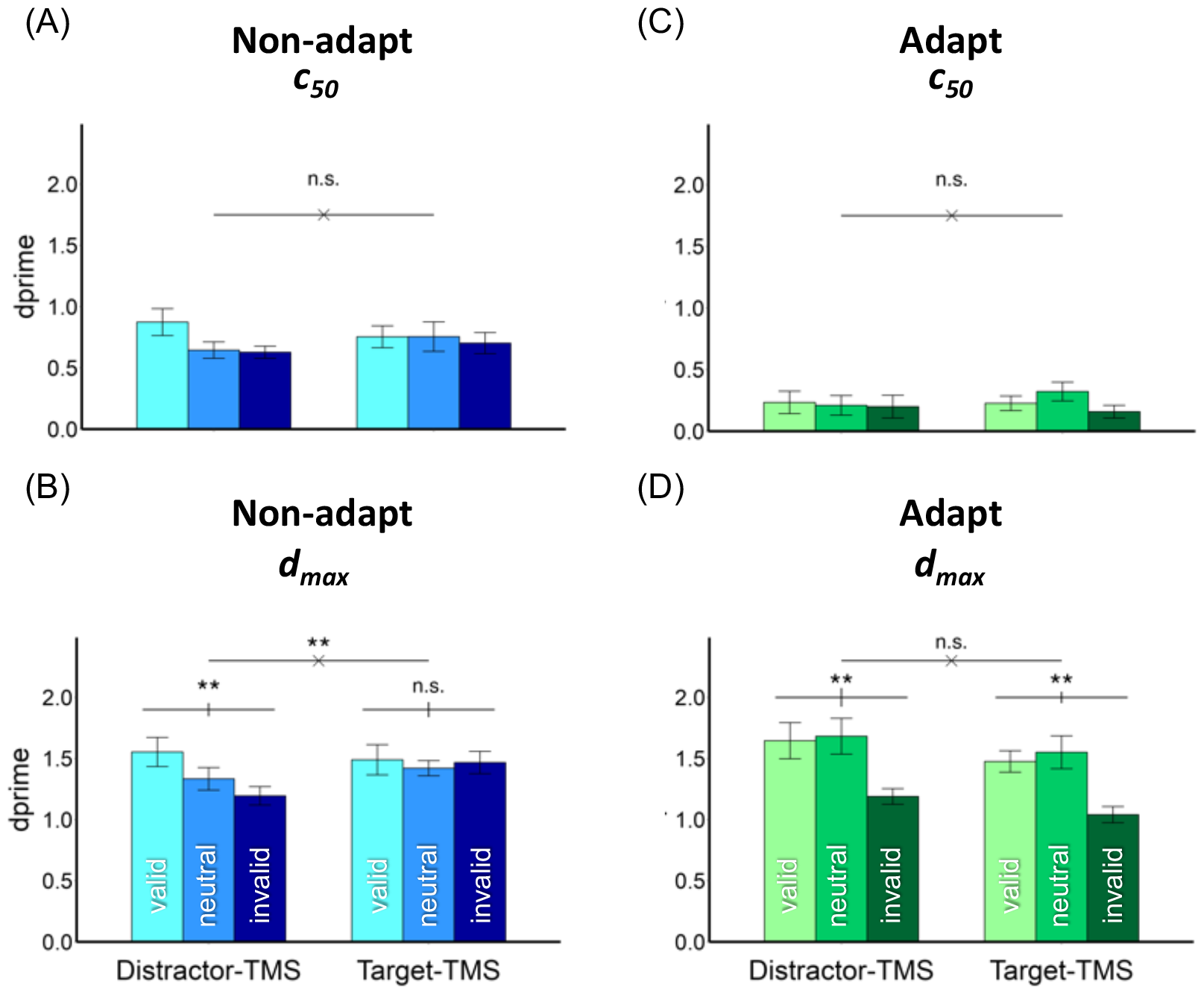
Performance as indexed by *d’* in the (A) non-adapt c_50_ condition, (B) non-adapt d_max_ condition, (C) adapt c_50_ condition, (D) adapt d_max_ condition. The error bars within the bar plots depict ±1 SEM (Cousineau corrected) of the condition. The error bars above the bar plots indicate ±1 SEM of the difference between the valid and invalid conditions. ** *p* < .01, * *p* < .05.

For d_max_ (**Figure 4B**), there was an interaction between attention and TMS [*F*(2,24)=4.23, *p*=.027, *η*^*2*^=0.26]; there was an attentional effect in the distracter-stimulated condition [*t*(12)=3.36, *p*=.006, *d*=0.93] but not in the target-stimulated condition [*t*(12)=0.18, *p*=.857]. When comparing the target-stimulated condition to the distractor-stimulated condition, the enhancement in the invalid condition (11 out of 13 observers) was more consistent across observers than the suppression in the valid condition (8 out of 13 observers). The finding that TMS eliminated the exogenous attentional effect is consistent with Fernández and Carrasco’s study^10^.

### The attentional effects under TMS with adaptation

For c_50_ (**Figure 4C**), there were neither main effects of attention [*F*(2,24)<1] nor TMS[*F*(1,12)=1.21, *p*=.293], nor was there an interaction [*F*(2,24)=1.09, *p*=.352]. For d_max_ (**Figure 4D**), there was a main effect of attention [*F*(2,24)=11.81, *p*<.001, *η*^*2*^=0.5], but no effect of TMS [*F*(1,12)=2.41, *p*=.146] or its interaction with attention [*F*(2,24)<1]. This result indicates that after adaptation, TMS did not influence the effect of exogenous attention on contrast sensitivity.

### Comparing the attentional effect with and without adaptation

To quantify the overall attentional effects, we calculated the difference in the valid *d’* and invalid *d’* values. **Figure 5** shows the comparison between the attentional effect with adaptation (y-axis) and without (x-axis) adaptation. In the distractor-stimulated condition, for c_50_ (**Figure 5A**), a 2-way ANOVA on adaptation and attention revealed a main effect of adaptation [*F*(1,12)=25.92, *p*<.001, *η*^*2*^=0.68], but not of attention [*F*(1,12)=3.08, *p*=.105], nor an interaction [*F*(1,12)=1.95, *p*=.187]. For d_max_ (**Figure 5B**), there was a main effect of attention [*F*(1,12)=14, *p*=.003, *η*^*2*^=0.54], but neither an effect of adaptation, nor an interaction [both *F*(1,12)<1]. Most individual data points in the scatterplot are along the diagonal.

**Figure 5.**
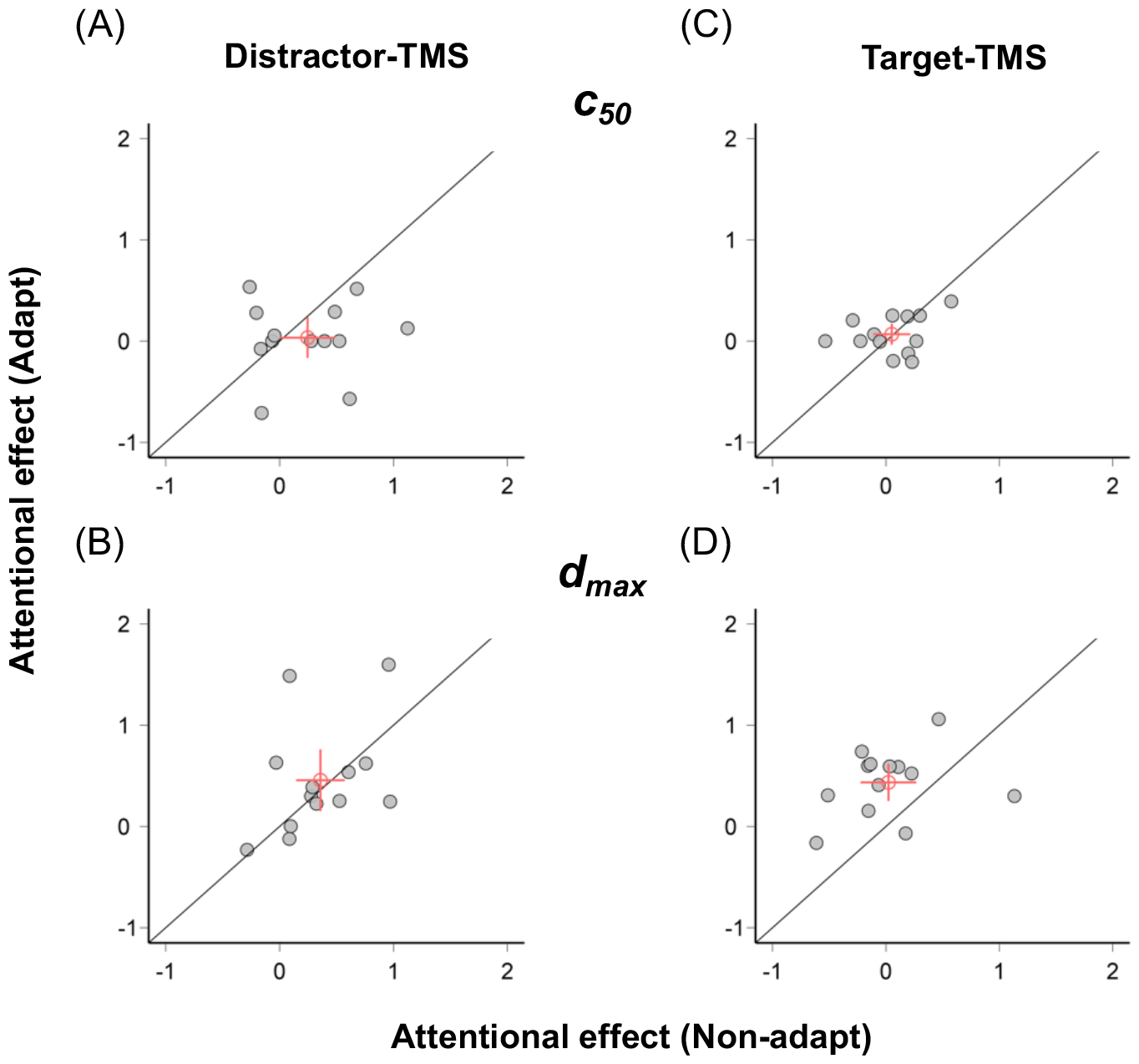
The attentional effect in the adapted (y-axis) and the non-adapted (x-axis) conditions for each observer in (A) distractor-TMS) c_50_, (B) distractor-TMS d_max_, (C) target-TMS c_50_, (D) target-TMS d_max_. The red circle indicates the average across observers and the error bars indicate ±1 SEM of the attentional effect.

In the target-stimulated condition, for c_50_ (**Figure 5C**), the 2-way ANOVA showed a main effect of adaptation [*F*(1,12)=60, *p*<.001, *η*^*2*^=0.83], but neither a main effect of attention [*F*(1,12)=1.25, *p*=.285] nor an interaction with adaptation [*F*(1,12)<1]. For d_max_ (**Figure 5D**), the 2-way ANOVA revealed an interaction [*F*(1,12)=9.63, *p*=.009, *η*^*2*^=0.45]; there was an attentional effect in the adapted condition [*t*(12)=4.73, *p*<.001, *d*=1.32] but not in the non-adapted condition [*t*(12)=0.18, *p=*.857]. Note that in the scatterplot, most points lie in the upper left diagonal. The scatterplot here provides supporting evidence across observers that after adaptation, TMS did not eliminate the effect of attention on performance.

## Discussion

Both adaptation and attention induce relatively short-term plastic changes, prioritizing relevant over irrelevant information, whether related to sensory history or behavioral relevance. These changes facilitate the visual system to manage limited resources and adjust to current environmental demands. In this study, we capitalized on previous findings showing that TMS^22,24^, exogenous covert attention^10^ and adaptation^18,19^, each alter brain state in early visual cortex. Here, to investigate the relation between adaptation and attention in the early visual cortex, we manipulated brain state through visual adaptation and attention in a psychophysical experiment while applying TMS.

In the distractor-stimulated condition, in which we expected no effect of TMS, we demonstrated (1) a contrast gain effect of adaptation (**Figure 3**), and (2) a response gain effect of exogenous attention (**Figure 4B**). These findings are consistent with those of an adaptation and attention study without neurostimulation^11^. The phosphene mapping procedure we used is similar to that in previous studies^10,20,21,25-29^. TMS-induced phosphenes are confined to the contralateral visual hemifield. Thus, the distractor-stimulated condition was an ideal control condition (see Methods), and the target-stimulated condition was the only one in which we expected TMS to disrupt target processing.

In the target-stimulated condition: (1) TMS to early visual cortex eliminated the exogenous attentional effect on contrast sensitivity (**Figure 4B**), replicating the study of Fernández and Carrasco^10^; and (2) adaptation eliminated the effect of TMS on attention (**Figure 4D**). These findings reveal that adaptation and attention interacted in the early visual cortex: by altering the brain state, adaptation enabled exogenous attention to exert its effects on performance and prevented it from being eliminated by TMS.

In the distractor-stimulated conditions, we observed typical adaptation and attentional effects, further indicating that it served as an ideal control condition. Specifically, adaptation shifted the CRF toward the 100%-contrast adaptor via contrast gain^6,11,18,23,30,31^ (**Figure 1A**). Additionally, exogenous attention multiplicatively enhanced neuronal firing rate as a function of contrast via response gain^7,11,32-34^ (**Figure 1B**). Specifically, in the distractor-stimulated condition, higher d_max_ was observed in the valid than invalid trials^2,7,10,11,33,35^ in both adapted and non-adapted conditions (**Figures 4B & 4D**). The fact that attention can modify perception after adaptation indicates that adaptation is not merely a by-product of neuronal fatigue; instead, attention can reset the system in a dynamic fashion^11,36^.

TMS induces a magnetic field to alter the neuronal activity^12-14,37-41^ and has been widely used to examine the causal or critical role of different brain regions for distinct perceptual and cognitive processes, such as motion perception^42,43^, face perception^44,45^, visual awareness^46,47^, multisensory information^48,49^, working memory^50^, and cognitive control^51,52^. The effects of TMS on human cortex are state-dependent^10,17-21,24,53-55^: TMS suppresses the excitatory activity, leading to a performance decrement, and the inhibitory activity (i.e., disinhibition), leading to a performance enhancement^10,18^.

In the current study, when attention was deployed to the target location, neural processing was enhanced at that location and depressed elsewhere. Thus, TMS eliminated the benefits in the valid condition while restoring the cost in the invalid condition. Specifically, in the target-stimulated condition without adaptation (**Figure 4B**). Thus, we replicated the extinction of exogenous attention’s effects on performance reported in Fernández and Carrasco^10^, confirming that the early visual cortex plays a causal role in the effect of exogenous attention on contrast sensitivity.

In the target-stimulated condition after adaptation, however, the effect of exogenous attention was not eliminated by TMS. TMS decreased the performance in the valid trials but did not improve performance in the invalid trials (**Figure 4D**). In the invalid adapted state at d_max_, performance was suppressed by both adaptation and the invalid cue, thus, the brain state might have been too suppressed for TMS to enable reactivation. These results suggest that the state-dependency effect of TMS has its limits: when brain activity is suppressed to a floor level, TMS could not reactivate it. Adaptation reduced contrast gain (i.e., the performance at c_50_^6,11,18,23,30^; **Figure 3**; however, adaptation can also suppress neural activity^56-58^ and behavior^59^ at higher contrasts. By using adaptation, attention and TMS simultaneously to alter the brain state, we provide evidence that adaptation and attention interact in the early visual cortex. This interaction could indicate that similar neuronal populations or similar mechanisms within different neural populations in early visual cortex underlie each effect.

By altering neural activity in V1/V2 with TMS, we reveal that the effect of exogenous attention, otherwise eliminated by TMS, was preserved by adaptation. This interaction between adaptation and exogenous attention provides a possible neural correlate for the psychophysical interaction in texture segmentation, where observers’ adaptation to high spatial frequencies eliminated the effect of exogenous attention at central locations^60^, as this task is also supported by the early visual cortex^9,61-64^. But adaptation and attention do not always interact behaviorally; they have independent effects on contrast sensitivity –observers’ adaptation to a Gabor did not modulate the beneficial effect of exogenous attention at the attended location and its cost at the unattended location^11^–and perceived speed –the effect of attention on perceived speed did not vary with adaptor speed^36^.

To conclude, we used a psychophysics-TMS protocol to investigate how adaptation modulates the effect of exogenous attention in the early visual cortex. We replicated both the typical contrast gain of adaptation and response gain of exogenous attention. Importantly, the extinction of exogenous attention effects on contrast sensitivity occurred when the target was disrupted by the TMS in the non-adaptation condition but not in the adaptation condition. Thus, adaptation shielded the attentional effect from disruption by TMS. This finding indicates that these two mechanisms, crucial for managing limited resources, interact at the initial level of cortical processing of visual information, possibly through similar neuronal populations.

## Limitation of the study

It has been reported that TMS impairs performance without adaptation, but restores performance after adaptation^18,19,55^. The following non-mutually exclusive factors may underlie why we did not observe these effects (**Figure 4C**): First, performance after adaptation at the c_50_ contrast was low. We titrated performance at ∼75% accuracy for c_50_ in the non-adapted, neutral condition and tested it in the adapted condition, where the accuracy for neutral trials was 51.15% in distractor-stimulated and 56.25% in target-stimulated condition, in line with 53% accuracy for the same condition in Perini et al.^18^. It is likely that neural activity may have reached a floor level and could not be reactivated. Second, the protocols differed: We stimulated one of the hemispheres during the task with two pulses and the intensity ranged from 49% to 65%. Perini et al.^18^ gave a single pulse at the center of the occipital pole with an intensity around 70%-80%, and the intensity of the TMS pulses can also influence the state-dependency of the TMS effect^16^. Third, instead of the no-TMS condition in Perini et al.^18^, we used a distractor-stimulated condition. Future studies could systematically examine how the protocol may influence the effects of TMS on cortical excitability and adaptation’s perceptual consequences.

## ACKNOWLEDGMENTS

This research was supported by NIH NEI R01-EY01 9693 to MC, NIH NINDS Grant F99-NS-120705 to AF, and the Ministry of Education in Taiwan to HHL. We thank current and former Carrasco Lab members, especially Ian Donovan, Laura Dugué, Nina Hanning and Shutian Xue, for their helpful comments.

## AUTHOR CONTRIBUTIONS

H.H.L., A.F. and M.C. designed research and interpreted the data; H.H.L. performed research, analyzed data, and drafted the paper; A.F. and M.C. guided and supervised the project; M.C. conceptualized the study, supervised and edited the writing of the paper, and provided funding.

## INCLUSION AND DIVERSITY

We worked to ensure gender balance as well as ethnic or other types of diversity in the recruitment of human subjects. While citing references scientifically relevant for this work, we also aimed to promote gender balance in our reference list.

## RESOURCE AVAILABILITY

### Lead contact

Further information and requests for resources should be directed to and will be fulfilled by the lead contact, Hsing-Hao Lee.

### Materials availability

This study did not generate new specimens or materials.

## Data and code availability

- The behavioral data has been uploaded the OSF database and are publicly available.
- This paper does not report original code.
- Any additional information required to reanalyze the data reported in this paper is available from the lead contact upon request.

## EXPERIMENTAL MODEL AND SUBJECT DETAILS

Thirteen observers (6 males, 7 females, including author HHL) participated in 4 experimental sessions, which is higher than the typical number of observers in previous TMS studies in visual perception^10,19,21,26,27,29,53^. We conducted bootstrapping based on our pilot study (n=3) to calculate the required sample size. With 10,000 iterations, bootstrapping results indicated that we would need 10 observers to reach a 3-way interaction among adaptation, attention, and TMS (power=90%). All observers were naïve to the purpose of the experiment and provided informed consent before participating in the experiment. All observers were free from neurological disorders and had normal or corrected-to-normal vision. This study followed the protocol of the safety guidelines for TMS research and was approved by the University Committee on Activities Involving Human Subjects at New York University.

## METHOD DETAILS

### Apparatus

The stimuli were presented on a gamma calibrated ViewPixx LCD monitor with 120 Hz refresh rate and 1920 × 1080 resolution. EyeLink 1000 (Eyelink SR) was used to monitor observers’ gaze (right eye) to make sure that observers were fixating at the fixation cross throughout the task and ensure that we were measuring a covert attentional effect. If observers moved their eyes (deviation > 1 dva) or blinked during the trial, the trial would stop and be repeated at the end of the block.

### Stimuli

The stimuli were generated using MATLAB (MathWorks, Natick, MA) and the Psychophysics toolbox^66,67^. The fixation cross consisted of two perpendicular lines (length=0.25 degree; width=0.06 degree) at the center of the screen. The Gabor patches (2 cpd) were presented on the left and right visual field, and the position was matched to the center of the reported phosphene by each observer [range: 4.24 – 14.34 dva away from the center]. The size of the Gabors were adjusted according to the cortical magnification factor^69^ : M = M_0_(1+0.42E+0.000055E^3^)^-1^. The attentional cues consisted of four solid black dots (0.1 dva wide), which surround the two Gabors (1 dva from the Gabor’s edge, 2 above/below, 2 left/right).

### Transcranial magnetic stimulation and phosphene mapping

The TMS pulses were given by a 70 mm figure-of-eight coil positioned at the occipital cortex with a Magstim Rapid Plus stimulator (3.5T) and triggered with MATLAB Arduino board. (Three observers received the pulses from an MCF-B70 coil with a MagVenture MagPro X100 stimulator instead). Stimulation intensity was the same throughout the experimental sessions for each observer and determined by the individual’s phosphene threshold (for 10 observers, 58%–65% of maximum Magstim stimulator output, mean = 61.3%, SD = 2.36%, and for the other 3 observers, 49%–58% of maximum MagVenture stimulator output, mean = 52%, SD = 5.2%, equipment was updated at NYU-TMS facility).

The phosphene mapping procedure was as the one used in previous studies^10,20,21,25-29^. Observers were seated 57 cm from the monitor in a dark room and were instructed to fixate at a dark-blue fixation at the center of a black background. A train of seven TMS pulses at 30 Hz and 65% intensity of the maximum output was applied on the occipital area of the scalp. Observers were instructed to draw the outline of the perceived phosphene on the screen using the mouse and the coil location was recorded accordingly. The center of the phosphene drawing was used as the coordinates of the Gabor’s location in the psychophysics task, where one Gabor was presented in the phosphene region (i.e., the stimulated region), and the other was presented in the symmetric region in the other hemifield (**Figure 2A**). The phosphene threshold was determined by two pulses spaced 50 ms apart at the same coil location. The intensity of the TMS pulse was adjusted accordingly until observers reported seeing phosphenes 50% of the time. The same phosphene mapping procedure was administered at the beginning of each session. The observer’s head was calibrated to match Brainsight software’s 3D head template, which ensured that the stimulation was given to the same location with the millimeter level of precision.

### Psychophysics-TMS task

After the phosphene mapping and before performing the psychophysics-TMS task in each session, we assessed the semi-saturation point and the asymptote of the CRF (i.e., the c_50_ and d_max_) by titrating the tilt angle and the contrast level for each observer (**Figures 1A & 1B**). They participated in thresholding tasks without adaptation and attention manipulation. We first conducted an adaptive staircase procedure to determine the tilt level (0.5° to 6° relative to vertical) that corresponds to approximately 75% orientation discrimination accuracy when the Gabor patches were presented with 80% contrast using the Palamedes toolbox^68^. Then, using the tilt level obtained from this tilt staircase task, we conducted a contrast staircase and varied the contrast of the Gabor from 5% to 30% to again achieved approximately 75% accuracy using the same toolbox to derive the semi-saturation point (c_50_; **Figure 1A**) of the CRF for each individual. The d_max_ contrast was fixed at 80% (**Figure 1B**) based on pilot data and previous studies with similar stimulus parameters^10,20,33,70^. The orientations of the left and right Gabors were independent of each other. We tested observers’ performance at these two contrasts across attention, adaptation, and TMS conditions to infer contrast gain and response gain mechanisms.

For the psychophysics-TMS task, **Figure 2B** shows the timeline and **Figure 2C** shows the experimental procedure. The adaptation and non-adaptation sessions were administered on different days to ensure that the adaptation effect did not carry-over to other conditions. The order of the adaptation and non-adaptation sessions was counterbalanced between observers.

In the adaptation blocks, observers were adapted to two cortically-magnified 100%-contrast Gabor patches (2cpd) on a mid-gray background flickering for 60 seconds in a counter phase manner at 10 Hz at the beginning of each block, followed by 2 seconds of top-up before each trial started (**Figure 2C**). This top-up was applied to ensure that the adaptation continued throughout the block. In the non-adaptation blocks, the procedure was the same but without the flickering Gabors; instead, a mid-gray screen was presented for 20 seconds followed by 2 seconds of blank at the beginning of each trial. Observers were instructed to fixate at the center and pay attention to the flickering Gabors during the adaptation phase.

After 400 ms of inter-stimulus interval (ISI), a valid, neutral, or invalid peripheral cue (40 ms) presented around a Gabor (1 dva away from the Gabor edge), followed by a 60 ms ISI, then two Gabor patches presenting at the center of placeholders (i.e., the center of the phosphene outline) on the left and right visual fields for 100 ms. Observers’ task was a two-alternative forced-choice orientation discrimination task (either counterclockwise or clockwise relative to the vertical) of the Gabor patch being indicated by the response cue, via button press. Note that the pre-cue was uninformative; in a random half of the trials, the response cue indicated the same location as precue (valid trials), while in the other half of the trials, the response cue indicated the other location (invalid trials).

During the target presentation, observers received two single pulses (separated by 50 ms) of TMS with the power at the sub-threshold level (**Figure 2B**). Like in previous studies^10,20,21,27^, we presented the two stimuli in the TMS stimulated region and its symmetric location in the other hemifield; the stimulus was presented for each observer according to their phosphene location (**Figure 2A**). In half of the trials, the response cue instructed observers to report the orientation of the stimulus at the stimulated region (contralateral to TMS; <u>target-stimulated</u>, which was equally likely to be a valid trial, invalid trial or neutral trial), and in the other half, the symmetric region (ipsilateral; <u>distractor-stimulated</u>, which was equally likely to be a valid trial, invalid trial or neutral trial). A feedback tone (400 Hz, 150 ms) was given after an incorrect response.

We used a lower intensity for the psychophysics-TMS task to ensure that no phosphenes were perceived during the main experimental task (conducted on mid-gray background). During the psychophysics-TMS task, if the stimulated region matched the response-cued region, it was a target-stimulated condition; otherwise, it was a distractor-stimulated condition (**Figure 2C**).

TMS over occipital cortex affects the contralateral hemifield^10^. Thus, the distractor-stimulated condition can be considered as a control condition (similar to a no-TMS condition), and the target-stimulated condition was the one in which TMS should disrupt target processing. Importantly, in our experimental design, observers could not know whether they were experiencing a valid or invalid cue trial (as the cue was uninformative) and whether they were in a target-stimulated or distractor-stimulated trial until the response cue appeared. Thus, the current experimental design eliminated the need for a sham condition, which can produce a different somatosensory experience than the sham^71,72^ and can bring expectation and placebo effects (see ^73^).

The whole experiment consisted of 4 sessions, and each session contained 10 blocks of 48 trials. Each observer completed 1920 trials in total, which included of 80 trials per condition (two different levels of contrast: c_50_ and d_max_; three attentional conditions: valid, neutral, and invalid; two adaptation conditions: adaptation and non-adaptation; two stimulated conditions: target-stimulated and distractor-stimulated (**Figure 2C**).

## QUANTIFICATION AND STATISTICAL ANALYSIS

Task performance indexed by *d’* [z(hit rate) – z(false alarm rate)] across conditions. The correct discrimination of clockwise trials were considered as hits and incorrect discrimination of counter-clockwise trials were considered as false alarms^10,20,32,70,74,75^.

Repeated-measures ANOVA along with effect size (*η*^*2*^) were computed in R^65^ and used to assess statistical significance. Partial *η*^*2*^ was provided for all F tests, where *η*^*2*^=0.01 indicates small effect, *η*^*2*^=0.06 indicates a medium effect, and *η*^*2*^=0.14 indicates a large effect. *Cohen’s d* was computed for each post-hoc *t*-test, where *d*=0.2 indicates a small effect, *d*=0.5 indicates a medium effect, and *d*=0.8 indicates a large effect^76^.

